# Identification of novel BDNF-specific corticostriatal circuitries

**DOI:** 10.1101/2021.08.26.457842

**Authors:** Yann Ehinger, Drishti Soneja, Khanhky Phamluong, Alexandra Salvi, Dorit Ron

## Abstract

BDNF is released from axon terminals originating in the cerebral cortex onto striatal neurons. Here, we characterized BDNF neurons in the corticostriatal circuitry. First, we utilized *BDNF*-Cre and Ribotag transgenic mouse lines to label BDNF-positive neurons in the cortex and detected *BDNF* expression in all the subregions of the prefrontal cortex (PFC). Next, we used a retrograde viral tracing strategy, in combination with *BDNF*-Cre knockin mice, to map the cortical outputs of BDNF neurons in the dorsomedial and dorsolateral striatum (DMS and DLS). We found that *BDNF-*expressing neurons located in the medial prefrontal cortex (mPFC) project mainly to the DMS, and those located in the primary and secondary motor cortices (M1 and M2) and agranular insular cortex (AI) project mainly to the DLS. In contrast, *BDNF-*expressing orbitofrontal cortical (OFC) neurons differentially target the dorsal striatum (DS) depending on their mediolateral and rostrocaudal location. Specifically, the DMS is mainly innervated by the medial and ventral part of the orbitofrontal cortex (MO and VO) whereas the DLS receives projections specifically from the lateral part of the OFC (LO). Together, our study uncovers previously unknown BDNF corticostriatal circuitries. These findings could have important implications for the role of BDNF signaling in corticostriatal pathways.

**Significance Statement:** BDNF is released in axons upon neuronal depolarization. Surprisingly, careful mapping of BDNF projecting neurons in the central nervous system (CNS) has not been conducted. Using retrograde viral strategies in combination with transgenic mice, we mapped out corticostriatal BDNF circuits. We found that, mPFC BDNF neurons project mainly to the DMS whereas the motor cortex and AI project to the DLS. BDNF neurons in the OFC are anatomically segregated. Whereas the DMS receives BDNF-positive projections from the VO, the DLS mainly receives BDNF-positive projections from the LO. Our findings could be important to the study of BDNF in corticostriatal circuitries.

## Introduction

Brain Derived Neurotrophic Factor (BDNF) is a member of the nerve growth factor family of neurotrophic factors, which is highly expressed in the CNS (Hofer et al., 1990; Yan et al., 1997; Kowianski et al., 2018). The majority of BDNF in neurons is stored in presynaptic dense core vesicles and is released upon neuronal depolarization (Dieni et al., 2012; Song et al., 2017). Once BDNF is released at axon terminals, it binds the receptor tyrosine kinase, tropomyosin-related kinase B (TrkB). Activation of TrkB stimulates extracellular regulated kinase 1/2 (ERK1/2), Protein kinase C (PKC) and/or phosphoinositide 3 kinase (PI3K) signaling cascades resulting in the initiation of transcriptional and translational machineries (Huang and Reichardt, 2003; Leal et al., 2014; Zagrebelsky et al., 2020). In the adult brain, BDNF plays a crucial role in synaptic and structural plasticity (Panja and Bramham, 2014; De Vincenti et al., 2019), as well as in learning and memory (Bekinschtein et al., 2014; Miranda et al., 2019).

BDNF is highly expressed in the cerebral cortex of both rodents and humans (Hofer et al., 1990; Timmusk et al., 1993; Conner et al., 1997). Studies in rodents suggest that BDNF in the cortex contributes to learning and memory paradigms (Miranda et al., 2019). For example, absence of BDNF in the prelimbic cortex (PrL) alters fear expression in mice indicating a role for consolidation and expression of learned fear (Choi et al., 2010). In the OFC, BDNF is critical for goal-directed decision making and in selecting actions based on their consequences (Gourley et al., 2013). In the motor cortex, BDNF contributes to motor learning (Andreska et al., 2020).

The cerebral cortex is also the major source of BDNF in the striatum (Altar et al., 1997; Baquet et al., 2004; Strand et al., 2007). BDNF released from cortical terminals, binds to, and activates its receptor, TrkB, in the striatum (Altar et al., 1997; Baydyuk and Xu, 2014). BDNF/TrkB signaling in the striatum has important cellular and behavioral roles (Besusso et al., 2013; Engeln et al., 2020). For example, TrkB signaling is required to control inhibition of locomotor behavior in enkephalin (ENK) positive medium spiny neurons (MSN) (Besusso et al., 2013), and Lobo and colleagues provided data to suggest that BDNF/TrkB signaling in dopamine D1 receptor expressing (D1) MSN plays a role in stereotypy behaviors (Engeln et al., 2020).

Finally, rodent studies have suggested that BDNF in corticostriatal circuitries is linked to addiction. For example, numerous studies investigated the role of BDNF in the PFC in relation with cocaine use (review (Pitts et al., 2016)). Specifically, cocaine exposure regulates BDNF signaling in the PFC and in turn BDNF influences the development and maintenance of cocaine-related behaviors (Lu et al., 2010; Gourley et al., 2013; Zhang et al., 2015; Pitts et al., 2018). In addition, activation of BDNF signaling in the DLS keeps alcohol intake in moderation (Jeanblanc et al., 2009; Jeanblanc et al., 2013), whereas malfunctioning of BDNF/TrkB signaling in the corticostriatal regions promotes compulsive heavy use of alcohol and other alcohol-mediated behaviors (Logrip et al., 2009; Darcq et al., 2015; Darcq et al., 2016; Warnault et al., 2016; Moffat et al., 2023).

As detailed above, BDNF is crucial for cortical and striatal functions, yet careful characterization of BDNF neurons in corticostriatal circuitry is lacking. Here, using a combination of transgenic mouse lines together with viral-mediated gene delivery approaches, we mapped out BDNF-expressing cortical neurons that project to the DMS and DLS.

### Methods Reagents

Mouse anti-NeuN antibody (MAB377) was obtained from Millipore (Billerica, MA). Rabbit anti-VGLUT1 antibody (VGT1-3) was purchased from Mab Technologies (Stone Mountain, GA). Chicken anti-GFP (A10262), donkey anti-mouse IgG AlexaFluor 594, anti-chicken AlexaFluor 488 and anti-rabbit IgG AlexaFluor 594 antibodies were purchased from Life Technologies (Grand Island, NY). Other common reagents were from Sigma Aldrich (St. Louis, MO) or Fisher Scientific (Pittsburgh, PA).

### Animals and Breeding

Male C57BL/6J mice (6-8 weeks old at time of purchase) were obtained from The Jackson Laboratory. Male *BDNF-*Cre knockin mice, which express Cre recombinase at the endogenous *BDNF* locus, were obtained from Zach Knight, UCSF (Tan et al., 2016). Ribotag mice (ROSA26CAGGFP-L10a), which express the ribosomal subunit RPL10a fused to EGFP (EGFP-L10a) in Cre-expressing cells (Zhou et al., 2013), were purchased from The Jackson Laboratory (B6;129S4-Gt (ROSA)26Sortm9(EGFP/Rpl10a)Amc/J). Ribotag mice were crossed with *BDNF-* Cre mice allowing EGFP-L10a expression in *BDNF-*expressing cells. Mouse genotype was determined by poly-chain reaction (PCR) analysis of tail DNA.

Mice were individually housed on paper-chip bedding (Teklad #7084), under a reverse 12-hour light-dark cycle (lights on 1000 to 2200 h). Temperature and humidity were kept constant at 22 ± 2°C, and relative humidity was maintained at 50 ± 5%. Mice were allowed access to food (Teklad Global Diet #2918) and tap water *ad libitum*. All animal procedures were approved by the university’s Institutional Animal Care and Use Committee and were conducted in agreement with the Association for Assessment and Accreditation of Laboratory Animal Care.

### Virus information

Recombinant adeno-associated virus (rAAV) retrograde EF1a Nuc-flox(mCherry)-EGFP (1 x 10^12^ vg/ml), which expresses nuclear-localized mCherry by default but switches to nuclear-localized EGFP expression in the presence of Cre (Back et al., 2019) (Addgene viral prep # 112677-AAVrg), and AAV1-pCAG-FLEX-EGFP, expressing EGFP in the presence of Cre (Addgene viral prep # 51502-AAV1), were purchased from Addgene.

### Stereotaxic surgery and viral infection

C57BL/6J (WT) or *BDNF*-Cre mice underwent stereotaxic surgery as described in Ehinger et al. (Ehinger et al., 2020). Specifically, mice were anesthetized by vaporized isoflurane and were placed in a digital stereotaxic frame (David Kopf Instruments, Tujunga, CA). A hole was drilled above the site of viral injection. The injector (stainless tubing, 33 gauges; Small Parts Incorporated, Logansport, IN) was slowly lowered into the target region. The injector was connected to Hamilton syringes (10 µl; 1701, Harvard Apparatus, Holliston, MA), and the infusion was controlled by an automatic pump at a rate of 0.1 µl/minute (Harvard Apparatus, Holliston, MA). The injector remained in place for an additional 10 minutes to allow the virus to diffuse and was then slowly removed. For experiments investigating BDNF-positive efferent projections to the DS, *BDNF*-Cre animals were unilaterally infused with 0.5 µl of retro-AAV-EF1aNuc-flox(mCherry)-EGFP targeting the DLS (anterior posterior (AP): +1.1, ML: ± 2.3, DV: −2.85, infusion at −2.8 from bregma) or the DMS (AP: +1.1, ML: ± 1.2, DV: −3, infusion at −2.95 from bregma). For experiments investigating BDNF positive efferent projections from the OFC, *BDNF*-Cre animals were unilaterally infused with 0.5 µl of AAV1-pCAG-FLEX-EGFP targeting the OFC (AP: +2.58, ML: ± 1.2, DV: −2.85, infusion at −2.8 from bregma).

### Immunohistochemistry, imaging and quantification

Following intraperitoneal (i.p.) administration of euthasol (200 mg/kg), mice were transcardially perfused with phosphate buffered saline, followed by 4% paraformaldehyde (PFA), pH 7.4. Brains were quickly removed post-perfusion and fixed for 24 hours in 4% PFA prior to cryoprotection in 30% sucrose solution for 3 days at 4°C. Brains were then sectioned to 30 µm by cryostat (CM3050, Leica, Buffalo Grove, IL), collected serially and stored at −80°C. PFA-fixed sections were permeabilized and blocked in PBS containing 0.3% Triton and 5% donkey serum for 4 hours at 4°C. Sections were then incubated for 18 hours at 4°C on an orbital shaker with the primary antibodies anti-NeuN (1/500), anti-GFP (1/1000) or anti-VGlut1 (1/1000) diluted in 3% bovine serum albumin (BSA) in PBS. Next, sections were washed in PBS and incubated for 4 hours at 4°C with Alexa Fluor 488-labeled donkey (1/500), Alexa Fluor 594-labeled donkey (1/500) antibodies in 3% BSA in PBS. After staining, sections were rinsed in PBS and were cover slipped using Prolong Gold mounting medium. Sections from rostral, rostrocaudal and caudal PFC were imaged on an Olympus Fluoview 3000 Confocal microscope (Olympus America, Center Valley PA) using manufacture recommended filter configurations, using the same parameters across mice and images. Captured images were used to quantify the number of fluorescent cells in subregions of rostral, rostrocaudal, and caudal PFC using FIJI ImageJ (NIH)(Schindelin et al., 2012). Specifically, the image of the PFC was precisely aligned to the anteroposterior (AP)-corresponding figure in the Paxinos atlas (Paxinos, 2004), using the BigWarp interface included in FIJI (J. A. Bogovic, 2016). By placing corresponding landmarks on the image and atlas, the image and the AP figure were overlayed with precision. Next, regions of interest (ROI) were traced on the overlayed image following the atlas delimitations using the polygon tool in FIJI. After adjusting the threshold (the same for each image), the number of positive neurons within an ROI was automatically quantified using the counter plugin in FIJI. To calculate the density of labeled neurons, the total number of labeled neurons in a region was divided by the surface of the region in mm^2^ using Fiji(Schindelin et al., 2012).

### Data Analysis

Graphpad Prism 9 was used for statistical analysis. D’Agostino–Pearson normality test was used to verify the normal distribution of variables. Data were analyzed using one-or two-way ANOVA where appropriate. One-way ANOVA was followed by Tukey’s multiple comparisons test when appropriate. For two-way ANOVAs, significant main effects or interactions were calculated, followed by Sidak’s multiple comparisons test. p value cutoff for statistical significance was set to 0.05.

## Results

### Experimental strategy for evaluating *BDNF*-expressing neurons distribution in the PFC

The cerebral cortex expresses high levels of *BDNF* message (Hofer et al., 1990; Timmusk et al., 1993), however, a careful analysis of *BDNF* containing neurons in the PFC has not been conducted. Therefore, we first assessed the distribution of *BDNF* expression in cortical neurons. To do so, we used a *BDNF*-Cre transgenic mouse line allowing Cre-recombinase expression only in *BDNF*-expressing cells (Tan et al., 2016), which was crossed with a Ribotag mouse line expressing GFP-fused ribosomal subunit RPL10 in the presence of Cre-recombinase (Zhou et al., 2013) (**Figure 1a**). The presence of GFP fused ribosomal subunit RPL10 (EGFP-L10a) enabled the visualization of *BDNF*-expressing cells.

**Figure 1:**
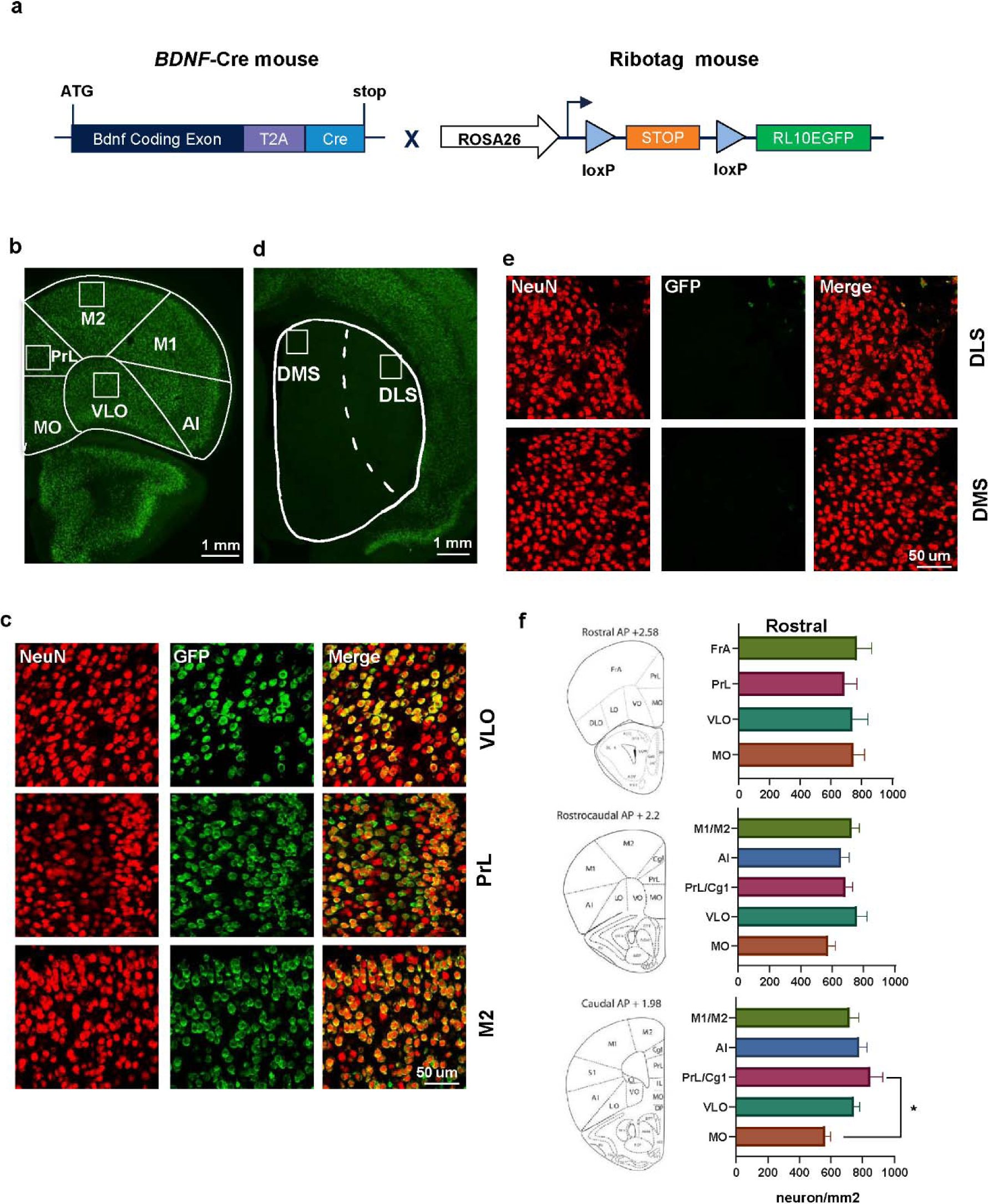
BDNF-positive neurons are detected in the PFC. (**a**) Experimental strategy for detection of *BDNF-*expressing neuron in the cortex. *BDNF*-Cre mice were crossed with Ribotag mice in which GFP-fused ribosomal subunit RPL10 is expressed in the presence of Cre recombinase, enabling the detection of *BDNF*-expressing neurons. (**b-c**) Representative images of cortical neurons. NeuN (red) and GFP (green) co-localization shows a high number of *BDNF*-expressing neurons in the mPFC, MC and OFC. (**d-e**) Representative images of the striatum. Neurons in the DMS and DLS do not express *BDNF*. (**f**) Topographical quantification of BDNF-positive neurons in the PFC along the rostrocaudal axis. One-way ANOVA followed by Tukey’s post hoc test, *p = 0.05, n = 5 mice. Scale bar is indicated on each panel. mPFC: medial prefrontal cortex, M1: primary motor cortex, M2: secondary motor cortex, PrL; prelimbic cortex, Cg1: Cingulate area 1, OFC: orbitofrontal cortex, VLO: ventrolateral OFC, MO: medial OFC, DMS: dorsomedial striatum, DLS: dorsolateral striatum, FrA: frontal association cortex.

### BDNF-positive neurons are detected in all prefrontal regions of the cortex

A large number of *BDNF-*expressing neurons were detected in the PFC, in M2 and the PrL as well as in the ventrolateral OFC (VLO) (**Figure 1b-c**). In contrast, we did not detect green fluorescence in the dorsal or ventral striatum (**Figure 1d-e**), which confirms previous data indicating that striatal neurons do not express *BDNF* (Baydyuk and Xu, 2014). We then quantified the number of *BDNF*-expressing neurons in the different PFC subregions and extended frontal regions (M1 and AI) along the rostrocaudal axis (**Figure 1f**). Rostral and rostrocaudal quantification revealed that all PFC subregions exhibit the same density of BDNF-positive neurons (rostral: 735.7±7.5 neurons/mm^2^, rostrocaudal: 749.3±11.3 neurons/mm^2^). We did not find significant differences in the number of BDNF-positive neurons between rostral subregions (one-way ANOVA, Df = 3, F (3, 16) = 0.7636, p = 0.5309, n = 5) and rostrocaudal (one-way ANOVA, Df = 4, F (4, 20) = 1.888, p = 0.152, n = 5). However, caudal quantification revealed a significantly lower density in BDNF-positive neurons in the MO compared to the PrLCingulate area 1(Cg1) (**Figure 1f**) (one-way ANOVA, Df = 4, F (4, 20) = 3.667, p = 0.0214; MO vs PrL/Cg1: p = 0.0122, n = 5).

### Experimental strategy for mapping of prefrontal BDNF-positive neurons projecting to the DS

The cerebral cortex is the major source of BDNF in the striatum (Altar, Cai et al. 1997, Baquet, Gorski et al. 2004). The striatum is divided into the ventral and dorsal striatum (Hunnicutt et al., 2016). We focused on the DS which is further divided into the lateral (DLS) and medial (DMS) regions (Hunnicutt et al., 2016), and assessed whether *BDNF*-expressing neurons in the PFC send projections to these two parts of the DS. To do so, we used a retrograde viral strategy and infected the anterior part of the DLS or the DMS of *BDNF*-Cre mice with a retrograde AAV-Nuc-flox-(mCherry)-EGFP viral construct (Back et al., 2019) (**Figure 2a**), which enables mCherry expression in BDNF-negative neurons and EGFP expression in BDNF-positive neurons (Back et al., 2019).

**Figure 2:**
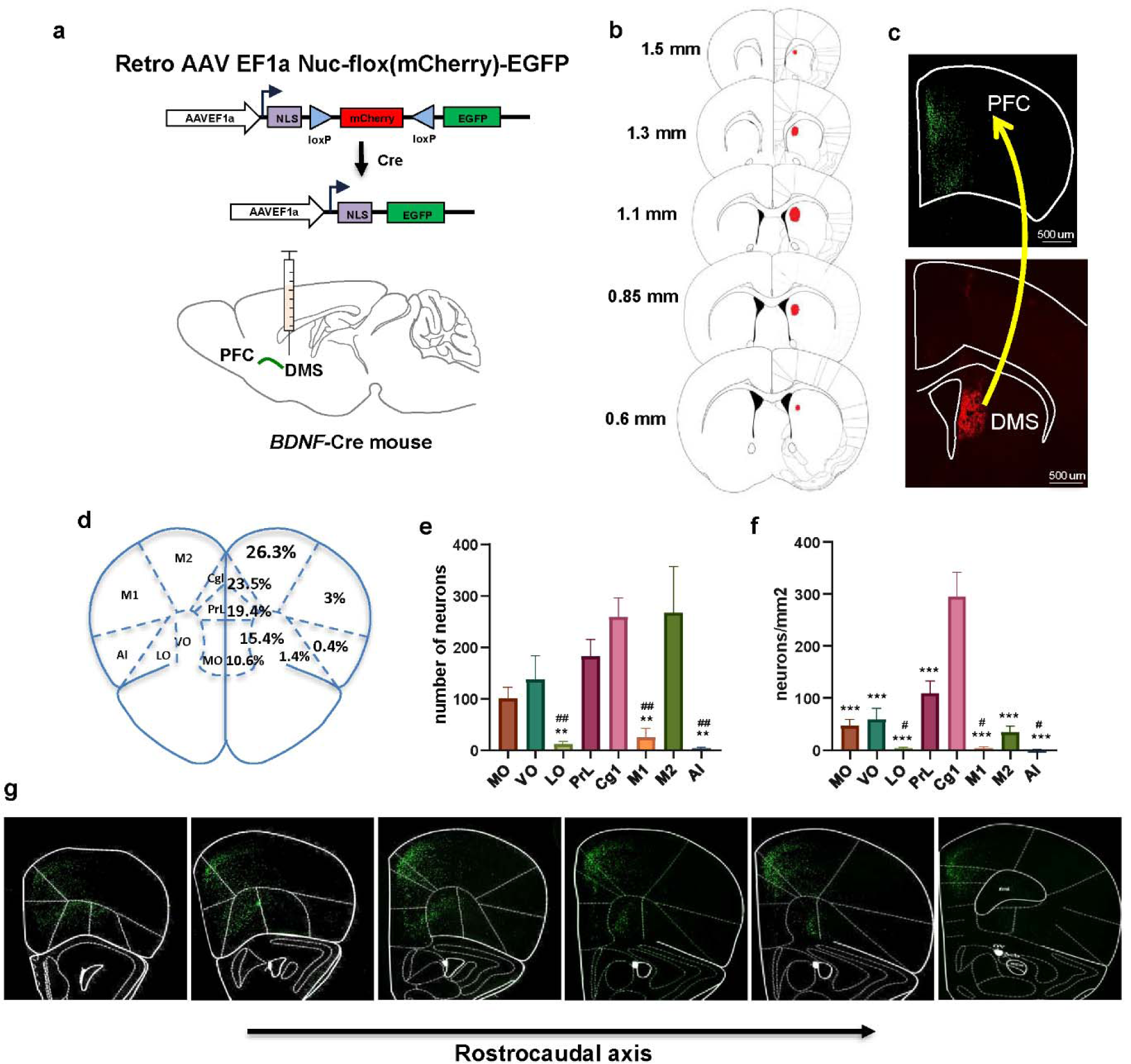
BDNF-positive neurons in the PFC project to the DMS. (**a**) Retrograde viral strategy to map *BDNF*-expressing cortical neurons projecting to the DMS. The DMS of *BDNF*-Cre mice was injected with a retrograde AAV EF1a Nuc-flox(mCherry)-EGFP viral construct enabling nuclear-localized mCherry to be expressed by default and to switch to nuclear-localized EGFP expression in the presence of Cre. (**b, h**) Retro AAV EF1a Nuc-flox(mCherry)-EGFP was infused in the DMS, and BDNF-positive neurons distribution was examined in the PFC. (**b**) Rostrocaudal distribution of the viral infection site. (**c**) *BDNF*-expressing neurons were identified by EGFP labeled nuclei and BDNF-negative neurons by mCherry nuclei. Lower panel shows BDNF-negative neurons (red) in the DMS. Upper panel shows *BDNF*-expressing neurons (green) in the PFC. (**d**) Percentage of total BDNF-positive projecting neurons per PFC subregion. (**e**) Number of BDNF-positive neurons projecting to the DMS per PFC subregion. (**f**) Density of BDNF-positive neurons projecting to the DMS per PFC subregion. (**g**) Representative images of PFC along the rostrocaudal axis. Contralateral BDNF-positive PFC projections to the DS were also quantified (**Extended Data Figure 2-1).** One-way ANOVA, followed by Tukey’s post hoc test. **^#^**p < 0.05, **^##^**p < 0.01 (PrL compared to other structures), **p < 0.01, ***p < 0.001 (Cg1 compared to other structures). n = 5 mice, scale bar is indicated on each panel. MO: medial OFC, VO: ventral OFC, LO: lateral OFC, PrL; prelimbic cortex, Cg1: Cingulate area 1, M1: primary motor cortex, M2: secondary motor cortex, AI: anterior insular cortex, DMS: dorsomedial striatum.

### PFC-to-DMS BDNF-positive circuits

First, we assessed whether *BDNF-*expressing cortical neurons project to the DMS by infecting the DMS of *BDNF*-Cre mice with the retrograde AAV-Nuc-flox-(mCherry)-EGFP (**Figure 2a-b**). As shown in **Figure 2c**, the DMS contains only cell nuclei labeled in red, corresponding to intrastriatal connections. The prefrontal regions show a high density of retrogradely EGFP labeled neurons (**Figure 2c and g**). Specifically, the mPFC including the PrL and Cg1 represents the major BDNF-positive output to the DMS, accounting for 42.9% of PFC-to-DMS projecting neurons (**Figure 2d-e**). In addition, 26.3% of the BDNF-positive projections to the DMS are coming from M2 (**Figure 2d-e**). The MO and VO represent respectively 10.6% and 15.4% of the BDNF-positive PFC-to-DMS projecting neurons (**Figure 2d-e**). Only 4.8% of total BDNF-positive neurons project from the LO, M1 and AI to the DMS (**Figure 2**).

### Density of *BDNF*-expressing neurons in PFC subregions that project to the DMS

To evaluate the density of the BDNF-positive PFC neurons projecting to the DMS within each PFC subregion, we analyzed the number of retrogradely labeled neurons per mm^2^ (**Figure 2f**). We found a significantly higher density of labeled BDNF-positive neurons in the Cg1 compared to the other regions, and in the PrL compared to LO, M1 and AI (**Figure 2f**) (one-way ANOVA, Df = 7, F (7, 32) = 22.72, p <0.0001; Cg1 vs other regions: p < 0.0001, PrL vs LO: p = 0.0206, PrL vs AI: p = 0.0159, PrL vs M1: p = 0.0204, n = 5). In addition, the BDNF-positive PFC neurons project bilaterally, but predominantly in an ipsilateral manner, and the topographic pattern of PFC *BDNF-*expressing neurons is similar in the contralateral hemisphere (**Extended Data Figure 2-1**).

### *BDNF*-expressing PFC neurons project to the DLS

Next, we assessed whether *BDNF*-expressing cortical neurons project to the DLS by infecting *BDNF*-Cre mice with the retrograde AAV-Nuc-flox-(mCherry)-EGFP (**Figure 3a-b**). Similar to the DMS, the DLS contained only red cell nuclei, corresponding to intrastriatal connections (**Figure 3c**). Absence of EGFP labeled neurons indicated once more a lack of *BDNF* message in the DLS. However, we found a high density of retrogradely EGFP labeled neurons in the prefrontal regions (**Figure 3c and g**). Specifically, the motor cortex (M1/M2), and the AI are the main regions that send BDNF-positive projection to the DLS, representing 80.9% and 12.6%, respectively, of the overall BDNF-positive projecting neurons (**Figure 3d-e**). Fewer BDNF-positive neurons in the OFC (MO, VO and LO) and mPFC (PrL and Cg1) project to the DLS, representing together only 6.4% of the overall BDNF-positive projecting neurons (**Figure 3d-e**).

**Figure 3:**
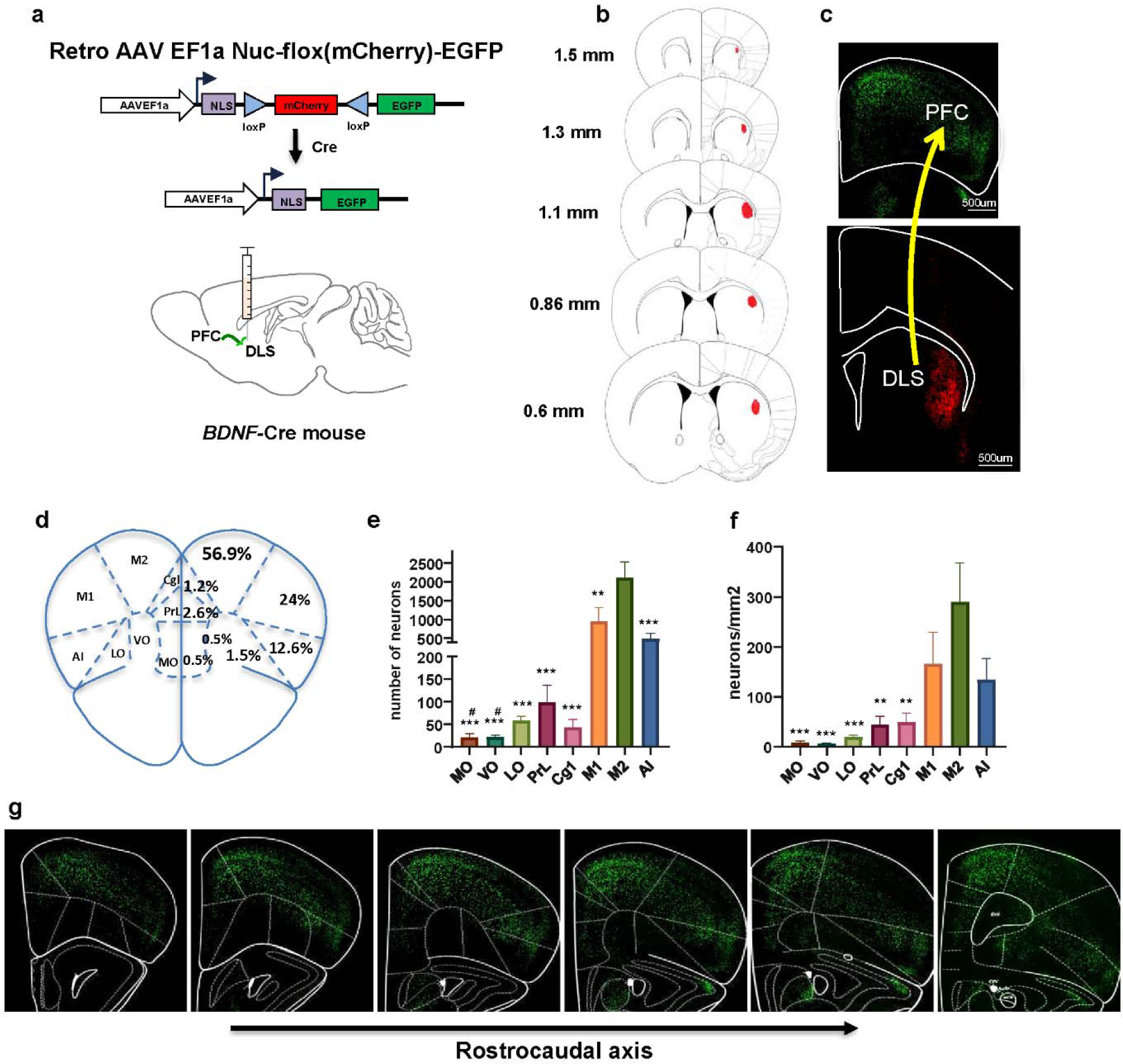
BDNF-positive neurons in the PFC project to the DLS. (**a**) Retrograde viral strategy to map *BDNF*-expressing cortical neurons projecting to the DLS. The DLS of *BDNF*-Cre mice was injected with a retrograde AAV EF1a Nuc-flox(mCherry)-EGFP viral construct enabling nuclear-localized mCherry to be expressed by default and to switch to nuclear-localized EGFP expression in the presence of Cre. (**b, h**) Retro AAV EF1a Nuc-flox(mCherry)-EGFP was infused in the DLS, and BDNF-positive neurons distribution was examined in the PFC. (**b**) Rostrocaudal distribution of the viral infection site. (**c**) BDNF-positive neurons were identified by EGFP labeled nuclei and BDNF-negative neurons by mCherry nuclei. Lower panel shows BDNF-negative neurons (red) in the DLS. Upper panel shows BDNF-positive neurons (green) in the PFC. (**d**) Percentage of total BDNF-positive projecting neurons per PFC subregion. (**e**) Number of BDNF-positive neurons projecting to the DLS per PFC subregion. (**f**) Density of BDNF-positive neurons projecting to the DLS per PFC subregion. (**g**) Representative images of PFC along the rostrocaudal axis. One-way ANOVA by Tukey’s post hoc test. *p<0.05, **p < 0.01, ***p < 0.001, M2 to other structures. n = 5 mice, scale bar is indicated on each panel. VO: ventral OFC, LO: lateral OFC, PrL; prelimbic cortex, Cg1: Cingulate area 1, M1: primary motor cortex, M2: secondary motor cortex, AI: anterior insular cortex, DLS: dorsolateral striatum.

### Density of PFC-to-DLS BDNF-positive projecting neurons in the PFC subregions

We next analyzed the density of retrogradely labeled BDNF-positive neurons in the PFC (**Figure 3f**). We observed that a higher number of labeled BDNF-positive neurons are located in M2 compared to the MO, VO, LO, PrL and Cg1 (one-way ANOVA, Df = 7, F (7, 32) = 6.412, p < 0.0001; M2 vs MO: p = 0.0004, M2 vs PrL: p = 0.0026, M2 vs VO: p = 0.0004, M2 vs LO: p = 0.0008, M2 vs Cg1: p = 0.0033, n = 5) (**Figure 3f**).

### Comparison of topographical distribution of BDNF-positive PFC to DLS and DMS projecting neurons along the rostrocaudal axis

We performed a more detailed comparison of PFC topographical patterns of DMS and DLS projecting neurons along the rostrocaudal axis (**Figure 4**). There was no significant difference between the PrL, VO, LO and MO retrogradely labeled BDNF-positive neurons at the rostral position (**Figure 4a**). Interestingly, rostrocaudal and caudal analysis revealed a specific AI-to-DLS circuit (**Figure 4b-c**). In contrast with the rostral position, retrogradely labeled BDNF*-*positive neurons in the PrL/Cg1 projecting to the DMS or DLS exhibit an opposite distribution, with fewer DLS projecting neurons and an increased number of DMS projecting neurons along the rostrocaudal axis (**Figure 4b-c**). We then focused on the OFC and compared the mediolateral distribution of BDNF-positive projecting neurons along the rostrocaudal axis (**Figure 4d**). We found that labeled *BDNF*-expressing neurons in the caudal LO are projecting significantly more to the DLS than the DMS, whereas labeled *BDNF*-expressing neurons in the rostral and rostrocaudal part of the VO are projecting significantly more to the DMS than the DLS (**Figure 4d**) (two-way ANOVA; interaction effect, F (8, 66) = 6.398, p < 0.0001, main effect of rostrocaudal axis, F (8, 66) = 3.177, p = 0.0042, main effect of OFC subregion, F (1.66) = 3.441, p = 0.0681, rostral VO: p = 0.0007, rostrocaudal VO: p = 0.011, caudal LO: p = 0.0028; n = 5).

**Figure 4:**
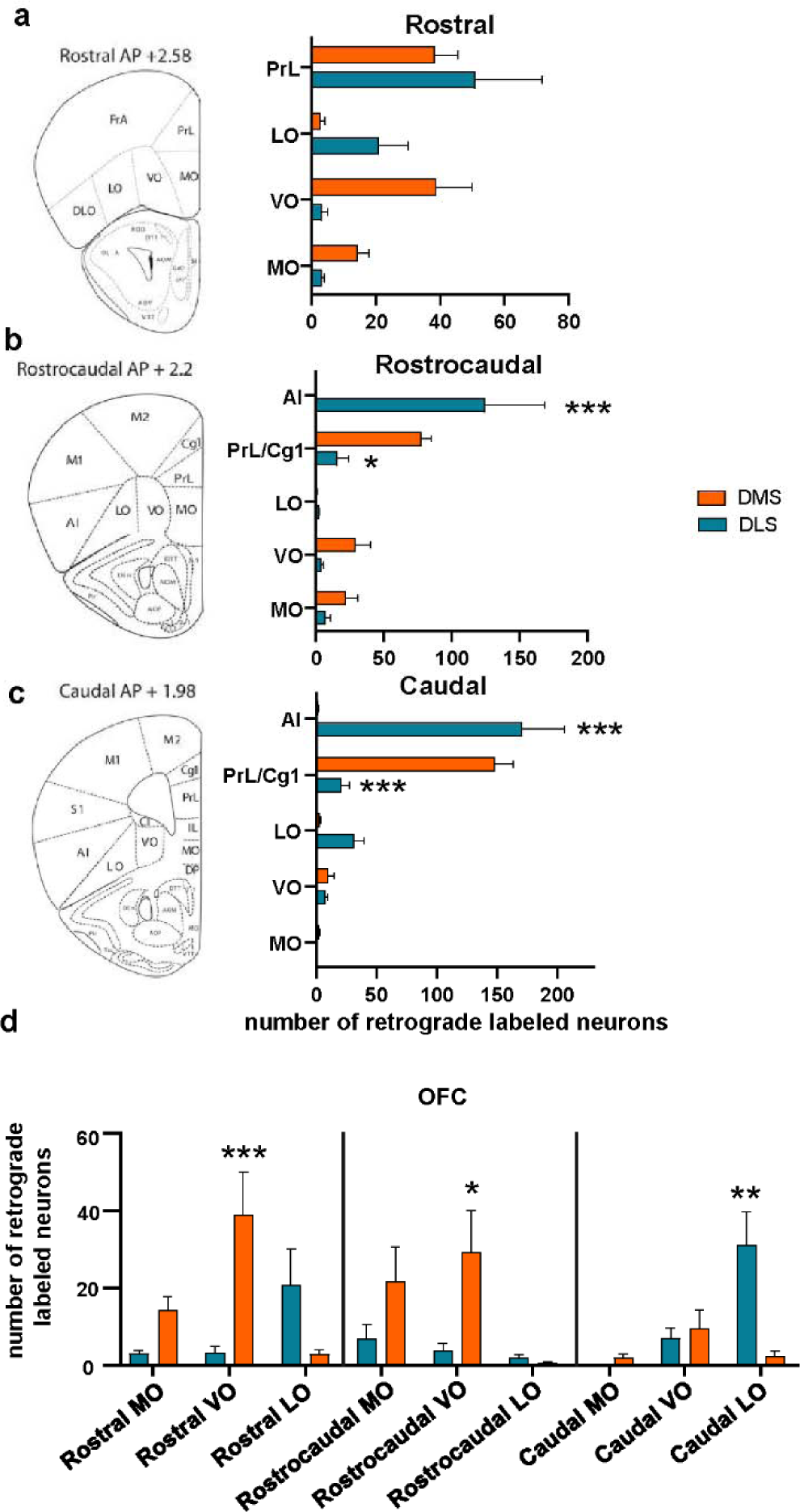
Comparison of topographical distribution of BDNF-positive PFC to DLS and DMS projecting neurons along the rostrocaudal axis. (**a-c**) Comparison of BDNF-positive PFC neurons projecting to the DMS versus DLS along the rostrocaudal axis. (**d**) Comparison of the distribution of BDNF-positive projecting neurons in the MO, VO and LO. Two-way ANOVA, followed by Sidak’s post hoc test. *p<0.05, **p < 0.01, ***p < 0.001, n = 5 mice. PFC: Prefrontal cortex, DLS: dorsolateral striatum, DMS: dorsomedial striatum, OFC: orbitofrontal cortex, MO: medial orbitofrontal cortex, VO: ventral orbitofrontal cortex, LO: lateral orbitofrontal cortex, PrL; prelimbic cortex, Cg1: Cingulate area 1, AI: anterior insular cortex.

### The DS receives inputs from BDNF-positive OFC neurons

Next, we used an anterograde viral strategy to confirm the presence of a BDNF-specific circuit between the OFC and DS. To do so, we infected the OFC of *BDNF*-Cre mice with an AAV-Flex-GFP virus allowing visualization of projections extended by BDNF-positive OFC neurons (**Figure 5a-c**). As shown in **Figure 5d**, BDNF-positive projections were localized in both the DLS and in the DMS.

**Figure 5:**
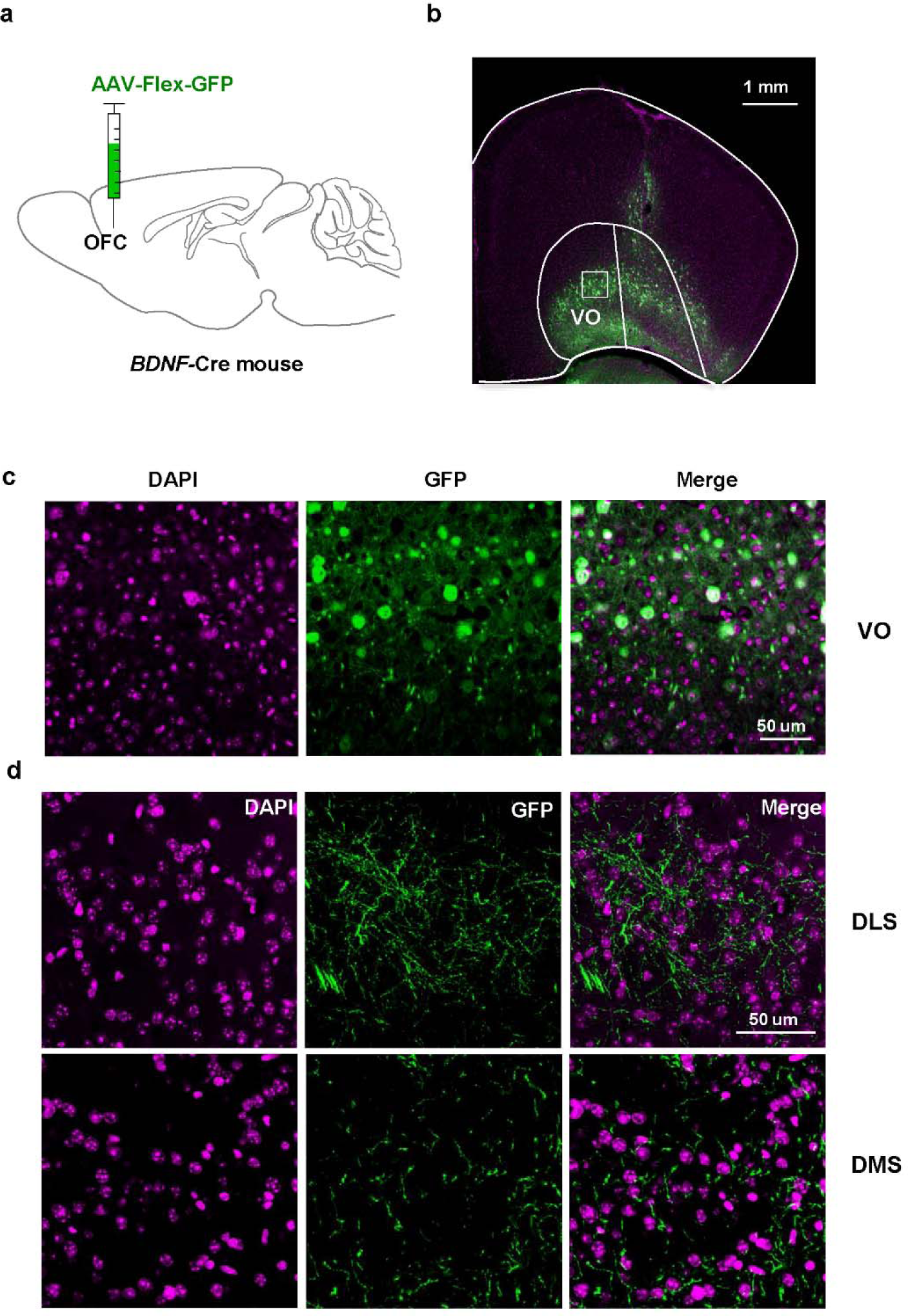
The DS receives projections from BDNF-positive OFC neurons. (**a-b**) Anterograde AAV1-Flex-GFP virus was injected in the OFC of *BDNF-*Cre mice allowing visualization of projections extended by BDNF-positive OFC neurons. (c-d) Representative images of the VO, DLS and DMS, showing GFP-positive projections localized in the DLS and in the DMS. DAPI staining is used to label nuclei. n = 2, scale bar is indicated on each panel. OFC: orbitofrontal cortex, DLS: dorsolateral striatum, DMS: dorsomedial striatum, DS: dorsal striatum, VO: ventral orbitofrontal cortex.

Finally, to determine whether the BDNF-positive OFC neurons form synapses with DLS neurons, we stained neurons with anti-vesicular glutamate transporter 1 (VGLUT1) antibodies labeling the cortical glutamatergic presynaptic compartment. GFP-positive projections and VGLUT1 were co-labeled in the DLS (**Figure 6**), suggesting that BDNF-positive glutamatergic neurons from the OFC extend projections and form synapses with neurons in the DLS.

**Figure 6:**
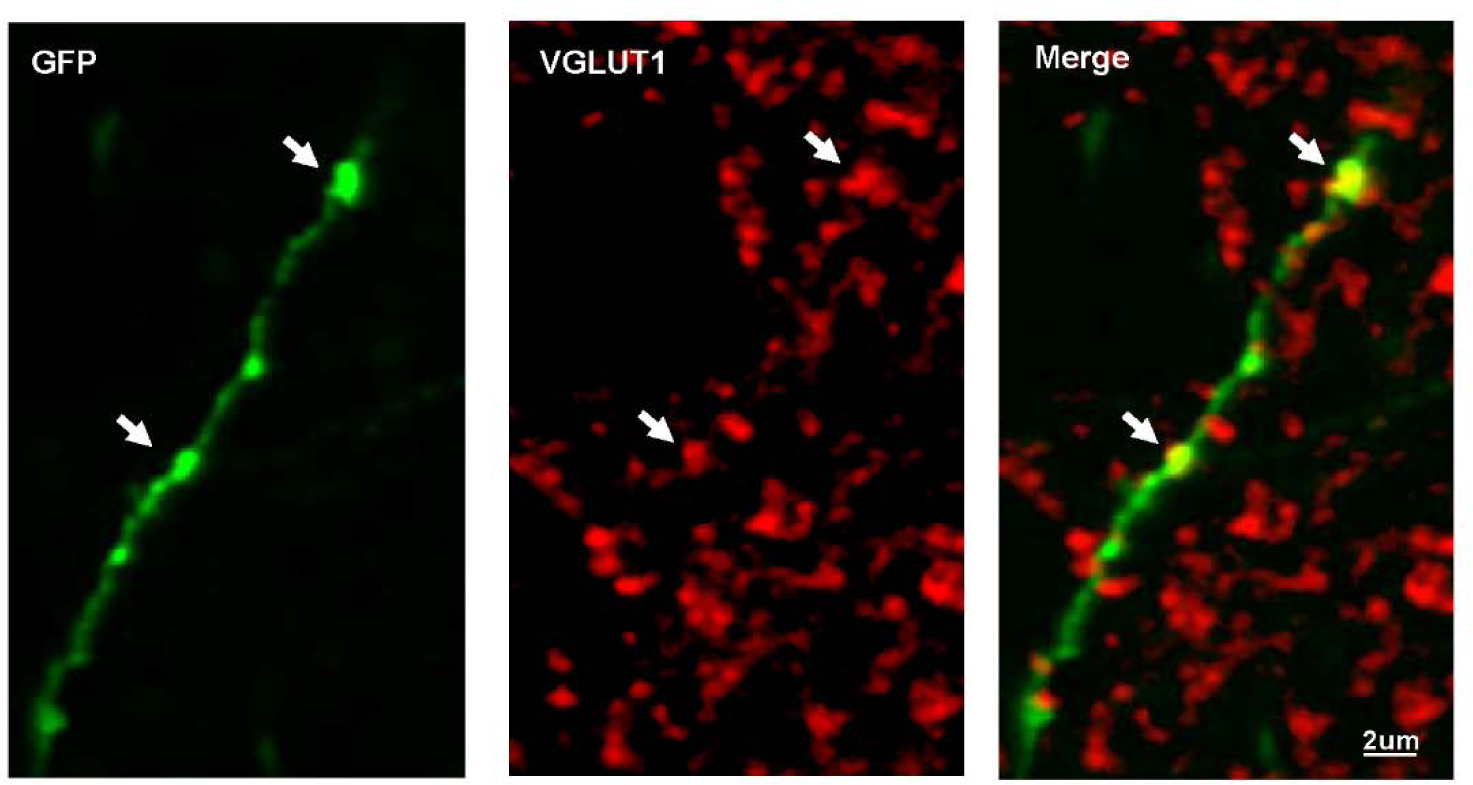
Glutamatergic BDNF-positive OFC neurons form synapses with DLS neurons. AAV-Flex-GFP virus was injected in the OFC of *BDNF-*Cre mice allowing visualization of projections in the DLS. GFP-positive projections in the DLS were stained for VGLUT1, a specific marker of cortical glutamatergic presynaptic compartment. Representative images of the DLS labeled with GFP in green and VGLUT1 in red. n = 2, scale bar is indicated on each panel. DLS: dorsolateral striatum, OFC: orbitofrontal cortex.

## Discussion

In the present study we mapped out cortical *BDNF*-expressing neurons projecting to the DLS and DMS. We found that *BDNF*-expressing neurons located in the mPFC project mainly to the DMS, whereas the motor cortex and the AI mainly project to the DLS. Interestingly, we observed an anatomical segregation along the rostrocaudal axis of OFC BDNF-positive neurons that project to the DLS and DMS. Thus, our data define novel BDNF neural circuits connecting the PFC to the DS.

### Characterization of BDNF neurons in the PFC and the DS

In 1990, Hofer et al. used *in situ* hybridization to characterize *BDNF* message in the brain and reported that *BDNF* mRNA is abundant in the mouse cerebral cortex (Hofer et al., 1990). In contrast, BDNF protein was detected in the striatum, even though *BDNF* mRNA was absent (Spires et al., 2004; Gharami et al., 2008). Altar et al. reported that BDNF protein found in the striatum is synthesized, anterogradely transported from the cell bodies located in the cerebral cortex and released presynaptically (Altar et al., 1997). Although more recent *BDNF* expression surveys have been conducted (Singer et al., 2018; Leschik et al., 2019; Wosnitzka et al., 2020), a careful mapping of *BDNF*-expressing neurons in cortical areas and their striatal projections has not been previously undertaken. Crossing a *BDNF*-Cre mouse line (Tan et al., 2016) with a Ribotag reporter line (Zhou et al., 2013) enabled labeling of *BDNF-*expressing neurons in the prefrontal cerebral cortex. Using this approach, which has superior accuracy compared to *in situ* hybridization and immunohistochemistry, we provide a definitive proof that a majority of neurons in prefrontal regions of the cortex express *BDNF*. Similar to what was previously reported (Hofer et al., 1990), we did not detect *BDNF*-expressing neurons in the striatum. Interestingly, although basal *BDNF* levels in the rodent striatum are negligible, a robust increase in *BDNF* message in the striatum is detected in response to behaviors such as voluntary alcohol intake (McGough et al., 2004; Jeanblanc et al., 2009) and cocaine administration (Liu et al., 2006), as well as in response to exercise (Marais et al., 2009) and stress (Miyanishi et al., 2021). Reconciling these potentially conflicting findings merits further investigation. One possibility is that these stimuli increase *BDNF* expression in cortical regions projecting to the striatum. Specifically, *BDNF* mRNA can be found in axon terminals (Lau et al., 2010), and recent studies show that presynaptic protein translation occurs within neuronal projections (Costa et al., 2019; Hafner et al., 2019). Thus, detectable *BDNF* in the striatum could be explained by the presence of *BDNF* mRNA within cortical neuron projections in the striatum.

### Characterization of corticostriatal BDNF circuitries

Our viral tracing strategy enabled us to explore BDNF-specific circuitries. For instance, we uncovered a specific AI-to-DLS BDNF circuit. Interestingly, Haggerty et al. recently found that stimulation of the AI-to-DLS circuit decreases alcohol binge drinking in male mice which in turn reshapes glutamatergic synapses in the DLS from AI inputs (Haggerty et al., 2022). In addition, we previously found that activation of BDNF/TrkB signaling in the DLS keeps alcohol intake in moderation (Jeanblanc et al., 2009; Jeanblanc et al., 2013). Thus, it is tempting to suggest a role for BDNF in AI-to-DLS synaptic adaptations that gate alcohol intake.

We observed that the mPFC exhibits a large number of BDNF-positive neurons projecting to the DMS. These results are in accordance with previous behavioral studies showing that the DMS receives projections from associative cortices such as the PrL (Friedman et al., 2015; Hart et al., 2018; Shipman et al., 2019; Vicente et al., 2020). We also found that a high number of BDNF-positive neurons in the motor cortex extends projections to the DLS. The motor cortex is known for innervating the DLS (Yin and Knowlton, 2006), and Andreska et al. recently reported that BDNF in this corticostriatal circuit is essential for motor learning (Andreska et al., 2020).

We also show herein that OFC BDNF-positive neurons project to the DS. Interestingly these projections are organized in a mediolateral manner. Specifically, the DMS receives projections from BDNF-positive neurons located in the MO and VO whereas BDNF-positive neurons projecting to the DLS are located specifically in the LO. It is known that the OFC projects to the DS (Gremel and Costa, 2013; Gremel et al., 2016; Zimmermann et al., 2017), and previous studies showed that the DMS receives input from the MO (Gourley et al., 2016; Green et al., 2020). The DMS plays an important role for goal directed behavior (Vandaele et al., 2019) and BDNF in the MO is essential to sustain goal-sensitive action in mice (Gourley et al., 2016). Specifically, Gourley et al. showed that *BDNF* knockdown in the MO decreases behavioral sensitivity to reinforcer devaluation. Thus, it is likely that this BDNF circuitry and perhaps BDNF released by MO neurons within the DMS regulate goal-directed action control.

We found that the majority of LO BDNF-positive neurons project to the DLS. The functional implication of this circuitry is unknow. Gourley and colleagues reported that BDNF in both VLO and MO participates in goal-directed behavior (Gourley et al., 2013; Gourley et al., 2016; Zimmermann et al., 2017), and Pitts et al. reported that inhibiting TrkB signaling in the DLS blocks habit formation (Pitts et al., 2018). In addition, mechanistic target of rapamycin complex 1 (mTORC1) in the VLO is involved in habitual alcohol seeking (Morisot et al., 2019). Putting these studies together, it is plausible that BDNF in the LO neurons projecting to the DLS plays a role in inflexible behavior.

Another possible role for BDNF in this corticostriatal circuit is to shape and modulate the synaptic output of striatal neurons. Using a two-neuron microcircuit approach in primary cortico-striatal neurons, Paraskevopoulou and colleagues showed that BDNF and glutamate co-released from cortical projections are required to modulate inhibitory synaptic transmission of striatal neurons (Paraskevopoulou et al., 2019). Therefore, co-release of BDNF with glutamate in the DLS and DMS by PFC neurons may modulate inhibitory sensitivity and dendritic morphology of striatal neurons in these PFC-DS circuits.

### BDNF-TrkB signaling in the DS

Most neurons in the striatum are MSN which are divided into two subpopulations, D1 MSN and dopamine D2 receptor (D2) expressing MSN (Gerfen et al., 1990). In the adult brain, both D1 and D2 MSN express TrkB receptors (Lobo et al., 2010). Engeln and colleagues showed that a subset of mice congenitally lacking TrkB receptor in D1-MSN in the DS exhibit repetitive circling behavior, suggesting a role of BDNF signaling in D1-MSN in this type of behavior (Engeln et al., 2020). Thus, it would be of great interest to map which dorsal striatal neuronal subtype receives inputs from cortical neuron. In addition, Nestler and colleagues provided data to suggest that activation of BDNF-TrkB signaling in D1 versus D2 MSN triggers opposite effects on cocaine and morphine-dependent rewarding behaviors in the nucleus accumbens (NAc) (Lobo et al., 2010; Koo et al., 2014). Additional studies are warranted to shed a light on the nature of cortical neurons projecting to the NAc. Furthermore, the DS is also organized into patch and matrix compartments which have distinct connectivity and genetic signature (Martin et al., 2019). As both of patch and matrix neurons express TrkB (Costantini et al., 1999), more studies are required to decipher the contribution of BDNF/TrkB signaling in subpopulation of DS neurons.

### Comparison between our findings and the literature

Using a retrograde viral tracing, we analyzed the spatial profiles of BDNF-positive projecting neurons in the PFC. We were able to confirm that these BDNF-positive circuits exhibit the same medial–lateral gradient of corticostriatal projections first reported in rats and primates (Selemon and Goldman-Rakic, 1988; Haber, 2003; Schilman et al., 2008). Although our data are generally consistent with the excitatory corticostriatal circuits described in the literature, several points might contribute to some discrepancies. For example, Balsters et al. compared corticostriatal circuits between human, non-human primates and mice and found significant differences in cortical projections to the anterior putamen and caudate body (Balsters et al., 2020). Many previous studies have been done in other animal models such as rat and primate (Berendse et al., 1992; Haber et al., 2006; Schilman et al., 2008; Hoover and Vertes, 2011; Mailly et al., 2013). Neuroanatomical differences exist across species and detailed explorations of similarities and differences between mice and other species are of importance. Furthermore, in contrast with previous literature (Berendse et al., 1992; Hoover and Vertes, 2011), in our comparison of the topographical distribution of BDNF-positive neurons, we observed VO-to-DMS and mPFC to-DLS BDNF-positive projecting neurons. An explanation to some of the discrepancies between corticostriatal circuit mapping studies is the experimental strategy used by us and others. Specifically, unlike the majority of studies that used anterograde strategies to map corticostriatal circuits (Berendse et al., 1992; Haber et al., 2006; Schilman et al., 2008; Hoover and Vertes, 2011; Mailly et al., 2013), we utilized a retrograde strategy in which a retrograde AAV was used (Tervo et al., 2016). Retrograde AAV is known to enter at the postsynaptic compartment of neurons and then be retrogradely transported to the cell body (Tervo et al., 2016). Thus, unlike an anterograde tracer which labels the entire fibers, we specifically labeled neurons projecting and forming synapses at the site of infection. However, Tervo et al. have provided good evidence to support the entry of AAV retrograde at axonal terminals (Tervo et al., 2016), additional studies will be necessary to fully rule out its ability to infect axons of passage. Although our anterograde tracing confirmed both the sparse and dense *BDNF*-expressing projections, anterograde tracing is not appropriate for detection of sparse projections.

Pan et al. reported tracing data consistent with our data using a fluorescent latex microsphere retrograde injection in mice (Pan et al., 2010). The authors reported an intense retrograde labeling in the PrL-IL and Cg1-M2 and no labeling in the LO-AI when injecting the retrograde tracer in the anterior DMS (aDMS). Pan et al. further found an intense retrograde labeling in the LO-AI and Cg1-M2 when injecting the fluorescent latex microsphere in the anterior DLS (aDLS). In addition, using a retrograde viral strategy, Green et al. also reported neurons located in the Cg1/PrL and IL/MO projecting to the aDMS and neurons located in the AI and M1/M2 projecting to the aDLS (Green et al., 2020).

It is important to note that our study focused specifically on the anterior part of the DS. Interestingly, Pan et al. found no retrograde labeling in the PrL, IL, MO, LO, AI, Cg1 and M2 when they injected the retrograde tracer in posterior part of the DS (pDS) (Pan et al., 2010). Since the pattern of projection in the striatum differs along the AP axis (Mailly et al., 2013; Hunnicutt et al., 2016), further work is needed to decipher PFC to pDS BDNF circuits in mice. In addition, using our BDNF-positive retrograde tracing strategy, the carefully mapped corticostrial circuities were restricted to the DMS and DLS and therefore lack dorsocentral inputs. Further tracing work is needed to describe the complete mediolateral PFC-to-DS circuitries.

In summary, in this study we mapped out *BDNF*-expressing neurons in the PFC and deciphered BDNF-specific corticostriatal circuits. Furthermore, the discovery of LO-to-DLS BDNF microcircuitry highlights the importance for deciphering the function of BDNF in the context of microcircuits composed by very localized neuronal ensembles.

---

**Extended Data Figure 2-1:**
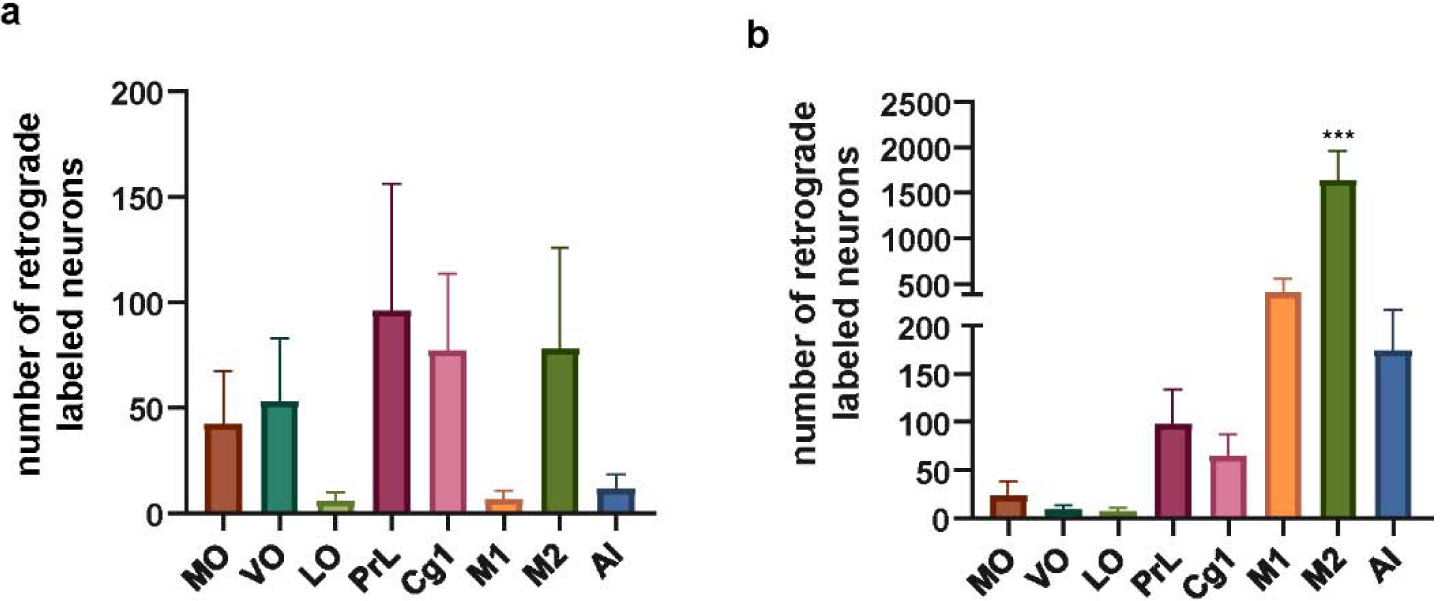
Contralateral BDNF-positive PFC projection to the DS. The DMS (**a**) or the DLS (**b**) of *BDNF-*Cre mice were injected with a retrograde AAV EF1a Nuc-flox(mCherry)-EGFP viral construct and retrogradely labeled neurons in the PFC were quantified. One-way ANOVA (F(7, 32) = 20.37; p<0.0001), followed by Tukey’s post hoc test. ***p < 0.001, * M2 compared to other structures. n = 5 mice. MO: medial OFC, VO: ventral OFC, LO: lateral OFC, PrL; prelimbic cortex, Cg1: Cingulate area 1, M1: primary motor cortex, M2: secondary motor cortex, AI: anterior insular cortex.

## References

Altar CA, Cai N, Bliven T, Juhasz M, Conner JM, Acheson AL, Lindsay RM, Wiegand SJ (1997) Anterograde transport of brain-derived neurotrophic factor and its role in the brain. Nature 389:856–860.

Andreska T, Rauskolb S, Schukraft N, Luningschror P, Sasi M, Signoret-Genest J, Behringer M, Blum R, Sauer M, Tovote P, Sendtner M (2020) Induction of BDNF Expression in Layer II/III and Layer V Neurons of the Motor Cortex Is Essential for Motor Learning. J Neurosci 40:6289–6308.

Back S et al. (2019) Neuron-Specific Genome Modification in the Adult Rat Brain Using CRISPR-Cas9 Transgenic Rats. Neuron 102:105–119 e108.

Balsters JH, Zerbi V, Sallet J, Wenderoth N, Mars RB (2020) Primate homologs of mouse cortico-striatal circuits. Elife 9.

Baquet ZC, Gorski JA, Jones KR (2004) Early striatal dendrite deficits followed by neuron loss with advanced age in the absence of anterograde cortical brain-derived neurotrophic factor. J Neurosci 24:4250–4258.

Baydyuk M, Xu B (2014) BDNF signaling and survival of striatal neurons. Front Cell Neurosci 8:254.

Bekinschtein P, Cammarota M, Medina JH (2014) BDNF and memory processing. Neuropharmacology 76 Pt C:677-683.

Berendse HW, Galis-de Graaf Y, Groenewegen HJ (1992) Topographical organization and relationship with ventral striatal compartments of prefrontal corticostriatal projections in the rat. J Comp Neurol 316:314–347.

Besusso D, Geibel M, Kramer D, Schneider T, Pendolino V, Picconi B, Calabresi P, Bannerman DM, Minichiello L (2013) BDNF-TrkB signaling in striatopallidal neurons controls inhibition of locomotor behavior. Nat Commun 4:2031.

Choi DC, Maguschak KA, Ye K, Jang S-W, Myers KM, Ressler KJ (2010) Prelimbic cortical BDNF is required for memory of learned fear but not extinction or innate fear. Proceedings of the National Academy of Sciences 107:2675–2680.

Conner JM, Lauterborn JC, Yan Q, Gall CM, Varon S (1997) Distribution of brain-derived neurotrophic factor (BDNF) protein and mRNA in the normal adult rat CNS: evidence for anterograde axonal transport. J Neurosci 17:2295–2313.

Costa RO, Martins H, Martins LF, Cwetsch AW, Mele M, Pedro JR, Tome D, Jeon NL, Cancedda L, Jaffrey SR, Almeida RD (2019) Synaptogenesis Stimulates a Proteasome-Mediated Ribosome Reduction in Axons. Cell Rep 28:864–876 e866.

Costantini LC, Feinstein SC, Radeke MJ, Snyder-Keller A (1999) Compartmental expression of trkB receptor protein in the developing striatum. Neuroscience 89:505–513.

Darcq E, Warnault V, Phamluong K, Besserer GM, Liu F, Ron D (2015) MicroRNA-30a-5p in the prefrontal cortex controls the transition from moderate to excessive alcohol consumption. Mol Psychiatry 20:1219–1231.

Darcq E, Morisot N, Phamluong K, Warnault V, Jeanblanc J, Longo FM, Massa SM, Ron D (2016) The Neurotrophic Factor Receptor p75 in the Rat Dorsolateral Striatum Drives Excessive Alcohol Drinking. J Neurosci 36:10116–10127.

De Vincenti AP, Rios AS, Paratcha G, Ledda F (2019) Mechanisms That Modulate and Diversify BDNF Functions: Implications for Hippocampal Synaptic Plasticity. Front Cell Neurosci 13:135.

Dieni S, Matsumoto T, Dekkers M, Rauskolb S, Ionescu MS, Deogracias R, Gundelfinger ED, Kojima M, Nestel S, Frotscher M, Barde YA (2012) BDNF and its pro-peptide are stored in presynaptic dense core vesicles in brain neurons. J Cell Biol 196:775–788.

Ehinger Y, Morisot N, Phamluong K, Sakhai SA, Soneja D, Adrover MF, Alvarez VA, Ron D (2020) cAMP-Fyn signaling in the dorsomedial striatum direct pathway drives excessive alcohol use. Neuropsychopharmacology.

Engeln M, Song Y, Chandra R, La A, Fox ME, Evans B, Turner MD, Thomas S, Francis TC, Hertzano R, Lobo MK (2020) Individual differences in stereotypy and neuron subtype translatome with TrkB deletion. Mol Psychiatry.

Friedman A, Homma D, Gibb LG, Amemori K, Rubin SJ, Hood AS, Riad MH, Graybiel AM (2015) A Corticostriatal Path Targeting Striosomes Controls Decision-Making under Conflict. Cell 161:1320–1333.

Gerfen CR, Engber TM, Mahan LC, Susel Z, Chase TN, Monsma FJ, Jr., Sibley DR (1990) D1 and D2 dopamine receptor-regulated gene expression of striatonigral and striatopallidal neurons. Science 250:1429–1432.

Gharami K, Xie Y, An JJ, Tonegawa S, Xu B (2008) Brain-derived neurotrophic factor over-expression in the forebrain ameliorates Huntington’s disease phenotypes in mice. J Neurochem 105:369–379.

Gourley SL, Zimmermann KS, Allen AG, Taylor JR (2016) The Medial Orbitofrontal Cortex Regulates Sensitivity to Outcome Value. J Neurosci 36:4600–4613.

Gourley SL, Olevska A, Zimmermann KS, Ressler KJ, Dileone RJ, Taylor JR (2013) The orbitofrontal cortex regulates outcome-based decision-making via the lateral striatum. Eur J Neurosci 38:2382–2388.

Green SM, Nathani S, Zimmerman J, Fireman D, Urs NM (2020) Retrograde Labeling Illuminates Distinct Topographical Organization of D1 and D2 Receptor-Positive Pyramidal Neurons in the Prefrontal Cortex of Mice. eNeuro 7.

Gremel CM, Costa RM (2013) Orbitofrontal and striatal circuits dynamically encode the shift between goal-directed and habitual actions. Nat Commun 4:2264.

Gremel CM, Chancey JH, Atwood BK, Luo G, Neve R, Ramakrishnan C, Deisseroth K, Lovinger DM, Costa RM (2016) Endocannabinoid Modulation of Orbitostriatal Circuits Gates Habit Formation. Neuron 90:1312–1324.

Haber SN (2003) The primate basal ganglia: parallel and integrative networks. J Chem Neuroanat 26:317–330.

Haber SN, Kim KS, Mailly P, Calzavara R (2006) Reward-related cortical inputs define a large striatal region in primates that interface with associative cortical connections, providing a substrate for incentive-based learning. J Neurosci 26:8368–8376.

Hafner AS, Donlin-Asp PG, Leitch B, Herzog E, Schuman EM (2019) Local protein synthesis is a ubiquitous feature of neuronal pre- and postsynaptic compartments. Science 364.

Haggerty DL, Munoz B, Pennington T, Viana Di Prisco G, Grecco GG, Atwood BK (2022) The role of anterior insular cortex inputs to dorsolateral striatum in binge alcohol drinking. Elife 11.

Hart G, Bradfield LA, Balleine BW (2018) Prefrontal Corticostriatal Disconnection Blocks the Acquisition of Goal-Directed Action. J Neurosci 38:1311–1322.

Hofer M, Pagliusi SR, Hohn A, Leibrock J, Barde YA (1990) Regional distribution of brain-derived neurotrophic factor mRNA in the adult mouse brain. EMBO J 9:2459–2464.

Hoover WB, Vertes RP (2011) Projections of the medial orbital and ventral orbital cortex in the rat. J Comp Neurol 519:3766–3801.

Huang EJ, Reichardt LF (2003) Trk receptors: roles in neuronal signal transduction. Annu Rev Biochem 72:609–642.

Hunnicutt BJ, Jongbloets BC, Birdsong WT, Gertz KJ, Zhong H, Mao T (2016) A comprehensive excitatory input map of the striatum reveals novel functional organization. Elife 5.

J. A. Bogovic PH, A. Wong and S. Saalfeld (2016) Robust registration of calcium images by learned contrast synthesis. IEEE 13th International Symposium on Biomedical Imaging (ISBI).

Jeanblanc J, Logrip ML, Janak PH, Ron D (2013) BDNF-mediated regulation of ethanol consumption requires the activation of the MAP kinase pathway and protein synthesis. Eur J Neurosci 37:607–612.

Jeanblanc J, He DY, Carnicella S, Kharazia V, Janak PH, Ron D (2009) Endogenous BDNF in the dorsolateral striatum gates alcohol drinking. J Neurosci 29:13494–13502.

Koo JW, Lobo MK, Chaudhury D, Labonte B, Friedman A, Heller E, Pena CJ, Han MH, Nestler EJ (2014) Loss of BDNF signaling in D1R-expressing NAc neurons enhances morphine reward by reducing GABA inhibition. Neuropsychopharmacology 39:2646–2653.

Kowianski P, Lietzau G, Czuba E, Waskow M, Steliga A, Morys J (2018) BDNF: A Key Factor with Multipotent Impact on Brain Signaling and Synaptic Plasticity. Cell Mol Neurobiol 38:579–593.

Lau AG, Irier HA, Gu J, Tian D, Ku L, Liu G, Xia M, Fritsch B, Zheng JQ, Dingledine R, Xu B, Lu B, Feng Y (2010) Distinct 3’UTRs differentially regulate activity-dependent translation of brain-derived neurotrophic factor (BDNF). Proc Natl Acad Sci U S A 107:15945–15950.

Leal G, Comprido D, Duarte CB (2014) BDNF-induced local protein synthesis and synaptic plasticity. Neuropharmacology 76 Pt C:639-656.

Leschik J, Eckenstaler R, Endres T, Munsch T, Edelmann E, Richter K, Kobler O, Fischer KD, Zuschratter W, Brigadski T, Lutz B, Lessmann V (2019) Prominent Postsynaptic and Dendritic Exocytosis of Endogenous BDNF Vesicles in BDNF-GFP Knock-in Mice. Mol Neurobiol 56:6833–6855.

Liu QR, Lu L, Zhu XG, Gong JP, Shaham Y, Uhl GR (2006) Rodent BDNF genes, novel promoters, novel splice variants, and regulation by cocaine. Brain Res 1067:1–12.

Lobo MK, Covington HE 3rd, Chaudhury D, Friedman AK, Sun H, Damez-Werno D, Dietz DM, Zaman S, Koo JW, Kennedy PJ, Mouzon E, Mogri M, Neve RL, Deisseroth K, Han MH, Nestler EJ (2010) Cell type-specific loss of BDNF signaling mimics optogenetic control of cocaine reward. Science 330:385-390.

Logrip ML, Janak PH, Ron D (2009) Escalating ethanol intake is associated with altered corticostriatal BDNF expression. J Neurochem 109:1459–1468.

Lu H, Cheng PL, Lim BK, Khoshnevisrad N, Poo MM (2010) Elevated BDNF after cocaine withdrawal facilitates LTP in medial prefrontal cortex by suppressing GABA inhibition. Neuron 67:821–833.

Mailly P, Aliane V, Groenewegen HJ, Haber SN, Deniau JM (2013) The rat prefrontostriatal system analyzed in 3D: evidence for multiple interacting functional units. J Neurosci 33:5718–5727.

Marais L, Hattingh SM, Stein DJ, Daniels WMU (2009) A proteomic analysis of the ventral hippocampus of rats subjected to maternal separation and escitalopram treatment. Metabolic Brain Disease 24:569–586.

Martin A, Calvigioni D, Tzortzi O, Fuzik J, Warnberg E, Meletis K (2019) A Spatiomolecular Map of the Striatum. Cell Rep 29:4320–4333 e4325.

McGough NN, He DY, Logrip ML, Jeanblanc J, Phamluong K, Luong K, Kharazia V, Janak PH, Ron D (2004) RACK1 and brain-derived neurotrophic factor: a homeostatic pathway that regulates alcohol addiction. J Neurosci 24:10542–10552.

Miranda M, Morici JF, Zanoni MB, Bekinschtein P (2019) Brain-Derived Neurotrophic Factor: A Key Molecule for Memory in the Healthy and the Pathological Brain. Front Cell Neurosci 13:363.

Miyanishi H, Muramatsu SI, Nitta A (2021) Striatal Shati/Nat8l-BDNF pathways determine the sensitivity to social defeat stress in mice through epigenetic regulation. Neuropsychopharmacology 46:1594–1605.

Moffat JJ, Sakhai SA, Hoisington ZW, Ehinger Y, Ron D (2023) The BDNF Val68Met polymorphism causes a sex specific alcohol preference over social interaction and also acute tolerance to the anxiolytic effects of alcohol, a phenotype driven by malfunction of BDNF in the ventral hippocampus of male mice. Psychopharmacology (Berl) 240:303–317.

Morisot N, Phamluong K, Ehinger Y, Berger AL, Moffat JJ, Ron D (2019) mTORC1 in the orbitofrontal cortex promotes habitual alcohol seeking. Elife 8.

Pan WX, Mao T, Dudman JT (2010) Inputs to the dorsal striatum of the mouse reflect the parallel circuit architecture of the forebrain. Front Neuroanat 4:147.

Panja D, Bramham CR (2014) BDNF mechanisms in late LTP formation: A synthesis and breakdown. Neuropharmacology 76 Pt C:664-676.

Paraskevopoulou F, Herman MA, Rosenmund C (2019) Glutamatergic Innervation onto Striatal Neurons Potentiates GABAergic Synaptic Output. J Neurosci 39:4448–4460.

Paxinos G, and Franklin, K. B. J. (2004) The Mouse Brain in Stereotaxic Coordinates. Elsevier Academic Press.

Pitts EG, Taylor JR, Gourley SL (2016) Prefrontal cortical BDNF: A regulatory key in cocaine- and food-reinforced behaviors. Neurobiol Dis 91:326–335.

Pitts EG, Li DC, Gourley SL (2018) Bidirectional coordination of actions and habits by TrkB in mice. Sci Rep 8:4495.

Schilman EA, Uylings HB, Galis-de Graaf Y, Joel D, Groenewegen HJ (2008) The orbital cortex in rats topographically projects to central parts of the caudate-putamen complex. Neurosci Lett 432:40–45.

Schindelin J, Arganda-Carreras I, Frise E, Kaynig V, Longair M, Pietzsch T, Preibisch S, Rueden C, Saalfeld S, Schmid B, Tinevez JY, White DJ, Hartenstein V, Eliceiri K, Tomancak P, Cardona A (2012) Fiji: an open-source platform for biological-image analysis. Nat Methods 9:676-682.

Selemon L, Goldman-Rakic P (1988) Common cortical and subcortical targets of the dorsolateral prefrontal and posterior parietal cortices in the rhesus monkey: evidence for a distributed neural network subserving spatially guided behavior. The Journal of Neuroscience 8:4049–4068.

Shipman ML, Johnson GC, Bouton ME, Green JT (2019) Chemogenetic Silencing of Prelimbic Cortex to Anterior Dorsomedial Striatum Projection Attenuates Operant Responding. eNeuro 6.

Singer W et al. (2018) BDNF-Live-Exon-Visualization (BLEV) Allows Differential Detection of BDNF Transcripts in vitro and in vivo. Front Mol Neurosci 11:325.

Song M, Martinowich K, Lee FS (2017) BDNF at the synapse: why location matters. Mol Psychiatry 22:1370–1375.

Spires TL, Grote HE, Varshney NK, Cordery PM, van Dellen A, Blakemore C, Hannan AJ (2004) Environmental enrichment rescues protein deficits in a mouse model of Huntington’s disease, indicating a possible disease mechanism. J Neurosci 24:2270–2276.

Strand AD, Baquet ZC, Aragaki AK, Holmans P, Yang L, Cleren C, Beal MF, Jones L, Kooperberg C, Olson JM, Jones KR (2007) Expression profiling of Huntington’s disease models suggests that brain-derived neurotrophic factor depletion plays a major role in striatal degeneration. J Neurosci 27:11758–11768.

Tan CL, Cooke EK, Leib DE, Lin YC, Daly GE, Zimmerman CA, Knight ZA (2016) Warm-Sensitive Neurons that Control Body Temperature. Cell 167:47–59 e15.

Tervo DG, Hwang BY, Viswanathan S, Gaj T, Lavzin M, Ritola KD, Lindo S, Michael S, Kuleshova E, Ojala D, Huang CC, Gerfen CR, Schiller J, Dudman JT, Hantman AW, Looger LL, Schaffer DV, Karpova AY (2016) A Designer AAV Variant Permits Efficient Retrograde Access to Projection Neurons. Neuron 92:372–382.

Timmusk T, Palm K, Metsis M, Reintam T, Paalme V, Saarma M, Persson H (1993) Multiple promoters direct tissue-specific expression of the rat BDNF gene. Neuron 10:475–489.

Vandaele Y, Mahajan NR, Ottenheimer DJ, Richard JM, Mysore SP, Janak PH (2019) Distinct recruitment of dorsomedial and dorsolateral striatum erodes with extended training. Elife 8.

Vicente AM, Martins GJ, Costa RM (2020) Cortico-basal ganglia circuits underlying dysfunctional control of motor behaviors in neuropsychiatric disorders. Curr Opin Genet Dev 65:151–159.

Warnault V, Darcq E, Morisot N, Phamluong K, Wilbrecht L, Massa SM, Longo FM, Ron D (2016) The BDNF Valine 68 to Methionine Polymorphism Increases Compulsive Alcohol Drinking in Mice That Is Reversed by Tropomyosin Receptor Kinase B Activation. Biol Psychiatry 79:463–473.

Wosnitzka E, Nan X, Nan J, Chacon-Fernandez P, Kussmaul L, Schuler M, Hengerer B, Barde YA (2020) A New Mouse Line Reporting the Translation of Brain-Derived Neurotrophic Factor Using Green Fluorescent Protein. eNeuro 7.

Yan Q, Rosenfeld RD, Matheson CR, Hawkins N, Lopez OT, Bennett L, Welcher AA (1997) Expression of brain-derived neurotrophic factor protein in the adult rat central nervous system. Neuroscience 78:431–448.

Yin HH, Knowlton BJ (2006) The role of the basal ganglia in habit formation. Nat Rev Neurosci 7:464–476.

Zagrebelsky M, Tacke C, Korte M (2020) BDNF signaling during the lifetime of dendritic spines. Cell Tissue Res 382:185-199.

Zhang Y, Zhu X, Huang C, Zhang X (2015) Molecular changes in the medial prefrontal cortex and nucleus accumbens are associated with blocking the behavioral sensitization to cocaine. Sci Rep 5:16172.

Zhou P, Zhang Y, Ma Q, Gu F, Day DS, He A, Zhou B, Li J, Stevens SM, Romo D, Pu WT (2013) Interrogating translational efficiency and lineage-specific transcriptomes using ribosome affinity purification. Proc Natl Acad Sci U S A 110:15395–15400.

Zimmermann KS, Yamin JA, Rainnie DG, Ressler KJ, Gourley SL (2017) Connections of the Mouse Orbitofrontal Cortex and Regulation of Goal-Directed Action Selection by Brain-Derived Neurotrophic Factor. Biol Psychiatry 81:366–377.

